# Distributed coding of space and valence in the medial temporal lobe

**DOI:** 10.1101/2024.11.16.623965

**Authors:** Ladan Yang, Catherine Mikkelsen

## Abstract

The hippocampus has been associated with spatial information processing (O’Keefe, 1976). However, it is also not clear whether such ensemble coding of spatial information extends to other brain regions in the medial temporal lobe. Hence, this study uses various classification techniques to attempt to decode spatial and valence information utilizing publicly available rodent single-unit data (Girardeau et al., 2017). We found that the spatial information is encoded sparsely and distributed among many neurons in the hippocampus. Additionally we found that both spatial and valence information could be decoded in the hippocampus, the amygdala, and the piriform cortex. We also compared the strengths and weaknesses of four different classifiers (LDA, KNN, SVM, NB) theoretically and in practice with this dataset.

## Introduction

### Encoding of spatial and valence information in the medial temporal lobe

Navigating through the environment and responding correctly to salient stimuli are important skills for animal survival. For example, they need to remember where they can find food and where they may be likely to encounter a predator. There is an extensive body of research connecting spatial navigation to its neural correlates. Recording the single units in the hippocampus, John O’Keefe and Jonathan Dostrovsky discovered place cells in the hippocampus that consistently fire when rats are in a specific location (1971). The Amygdala has been shown to be important for valence processing (Adolphs et al., 1994, LeDoux, 2007, Maren, 2001). Similar to the place cells, researchers found fear neurons in the amygdala that showed preferential firing to the conditioned stimulus (CS) after fear conditioning and reversed after extinction of the CS (Herry et al., 2008). The piriform cortex is typically associated with odor processing (Bekkers & Suzuki, 2013). This body of work has supported the idea of functional specialization, in which each area of the brain has a unique role to play in cognitive function. We sought to investigate the possibility of the role of distributed coding in the medial temporal lobe, in which brain areas might encode information beyond their “preferred domain”.

### Distributed coding of space and valence

While early work focused on specific brain regions, or subgroups of cells within aregion, that show distinct activation patterns in response to the stimuli of interest, more and more recent studies show that the information is encoded in the pattern across many cells or regions. What’s more, compared to the locationist view, distributed coding represents the idea that while somewhat specialized, brain areas may still code information about a variety of task dimensions (Rigotti et al., 2013, Eichenbaum, 2017). Recent evidence suggests that the hippocampus may implement a sparsely distributed coding method for episodic memories and spatial encoding such that it is selective to multiple dimensions of experience (Rigotti et al., 2013, Wixted et al., 2014, Stefanini et al., 2020). A recent study conducted by Poo et al. (2021), shows that, while the piriform cortex is known for coding odors, spatial locations can be decoded from activity in the posterior piriform cortex in rats during an odor-cued spatial choice task. It has also been found that the amygdala and piriform cortex employ a distributed coding method for valence and odor coding separately (Blazing & Franks, 2020). Thus, the first goal of this study is to test whether spatial location and salient cues can be decoded from single neuron firing patterns in the medial temporal lobe, specifically, in the hippocampus, central amygdala, basolateral amygdala, and piriform cortex.

### Classifiers

The finding of distributed coding is supported by the rising application of machine learning and classification techniques in neuroscience data. These methods enable researchers to classify the related event based on the pattern of a group of neurons, rather than focusing on a single neuron at a time.

Implementing these methods allow researchers to investigate mechanisms of behavior that may be more difficult to detect with univariate analysis (Haxby et al., 2001). Typically, data is first organized into two dimensions: samples and features. For example, firing rate of each neuron may constitute a feature, and the firing rate within a single trial could be a sample. Each of the samples is then matched with a corresponding label of interest, such as the spatial location of the rat. These data are then used to train the classifier so that the correspondence between the brain activity pattern and the stimuli of interest is learned. The classifiers can be used to predict the labels of a new dataset. Such population analysis is widely used in neuroscience, across diverse applications (Haynes & Rees, 2006, Alanazi, 2022, Park et al., 2021, Sarmashghi et al., 2022, Astrand et al., 2014). This method not only allows researchers to increase the accuracy of clinical prediction, but also helps elucidate the neural mechanisms of a behavior of interest.

However, with the popularity of new methods also comes the problem of selecting the best tool for a given question. It is worth noting that there is no algorithm that is best for every dataset and situation, a theorem term as “No Free Lunch” in ML (Wolpert & Macready, 1997). Thus, it is important to understand the differences between classifiers and select the optimal classifiers based on the data type and question in mind. There are only few studies comparing the classifiers using single cell recording for biological state prediction (Glaser et al., 2020), despite extensive work looking at classifier methods in other fields (Haynes & Rees, 2006, Misaki et al., 2010, Sarmashghi et al., 2022). Hence, the secondary goal of this study is to compare the performance of different classifiers in decoding behavior of interest using single unit recording data.

There are two main types of models in machine learning studies, discriminative and generative. Discriminative models compute only the posterior probability, the chance of belonging to a class given the data. The generative models, however, also compute the joint probability, the chance that two events both happen (Bishop, 2006). This distinction means generative models are more powerful and resources demanding. Thus, it can be wasteful of computational resources to use generative models when only the classification decisions are needed. However, since the marginal density of the data point is estimated in the generative model, it can increase the accuracy of classification in the minority class and novel data points (Bishop, 1993). Additionally, when selecting a model one must decide whether to use a linear classifier, which assumes a linear relationship between features and labels, or a non-linear classifiers, which does not assume linearity. We selected Linear Discriminant Analysis (LDA) and Gaussian Naive Bayes (GNB) as generative model representatives and linear Support Vector Machine (lin-SVM) and K-Nearest-Neighbor (KNN) as discriminative model representatives. Additionally, LDA and SVM are linear models, and NB and KNN are non-linear models. The choices are made based on their high performance displayed in previous research and their popularity (Misaki et al., 2010, Mandelkow et al., 2016, Mitchell et al., 2003, Yang et al., 2014, Shinde et al., 2023). Next, we will compare these classifiers, including their mechanism, assumptions, strengths and weaknesses (summarized in table 1).

**Table 1.**
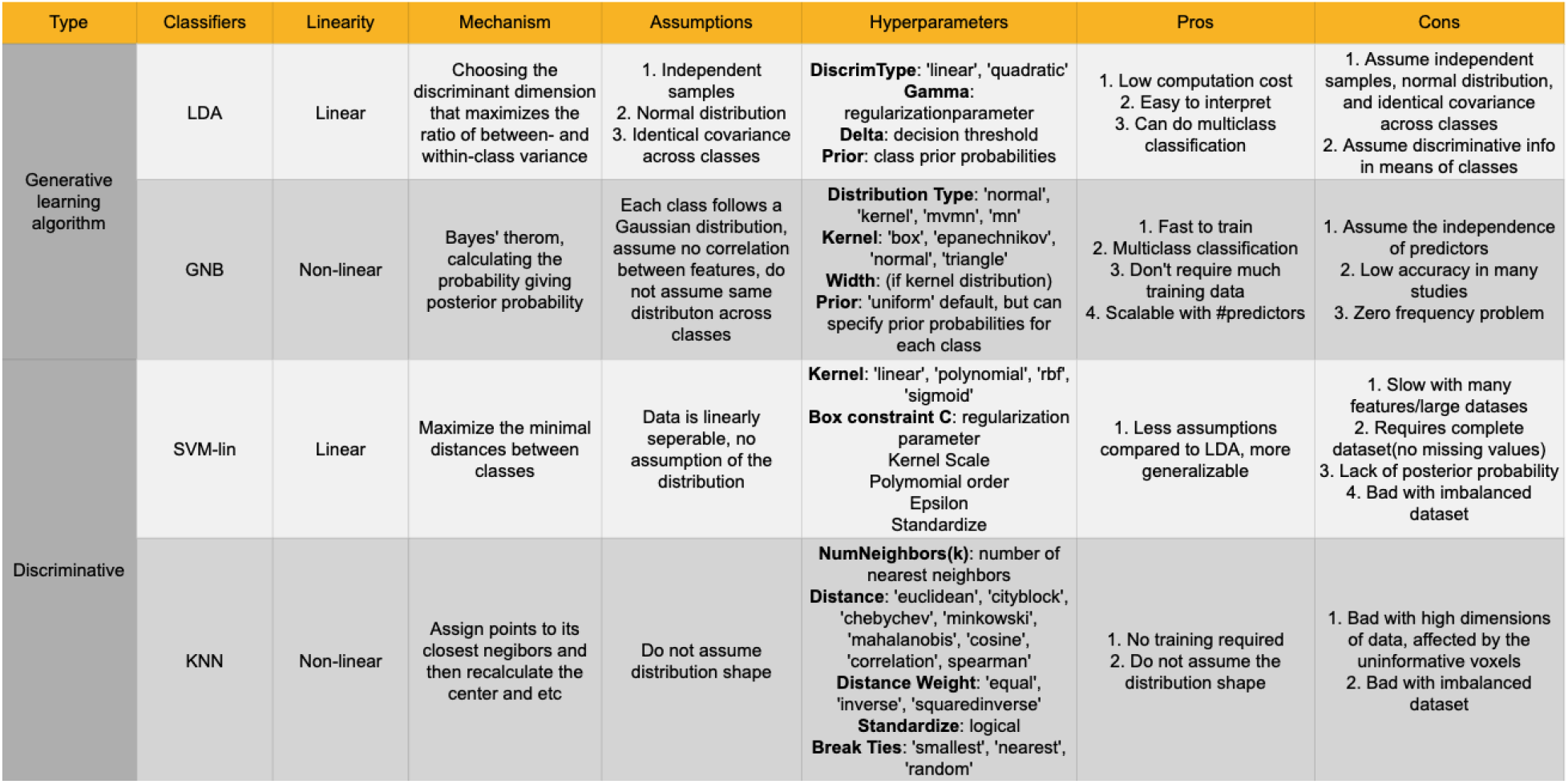
Comparisons of different classifiers.

### Linear Discriminant Analysis (LDA)

LDA is a classifier that linearly combines each features’ data to predict the outcome of interest. It assumes that samples are independent of each other and normally distributed. It also assumes an identical covariance matrix across classes, meaning that all classes share the same distribution, variance and correlation within features and only differ in means (Fisher, 1936). As one of the simplest classifiers, LDA is fast to train and easy to interpret, with the weights representing the relative importance of each feature (Duda et al., 2000, Bishop, 2007). However, because of these assumptions, LDA performs poorly when there are outliers, or when there are subclasses/clusters within classes, i.e. the data is multimodal (Qu & Pei, 2024). It is also not the best choice if the sample size is smaller than the dimensions of the data, which will result in overfitting.

### Gaussian Naive Bayes (GNB)

GNB is a nonlinear classifier that assumes each class follows a Gaussian distribution and that there is no correlation between features (Mitchell, 1997, Mitchell et al., 2004). It is fast and easy to train and it handles high dimensional data well. What’s more, it accounts for the probability of occurrence for each class. Yet, such dependence on prior probability of each class also makes it susceptible to the zero frequency problem, in which the lack of a class or feature in training will lead to zero frequency-based probability estimate. In the same vein, GNB is also susceptible to poor performance when the data is imbalanced (Kim & Lee, 2023).Table 1. Comparisons of different classifiers

### Support Vector Machine (SVM)

SVM projects the data into a hyperplane, a plane in the high dimensional space, that has the best separability between classes and uses the points near the hyperplane as a support vector for classification. One of the advantages is its many kernels which allows it to encompass simple linear and more complicated nonlinear models. Assuming linear separability only, SVM-lin is generalizable to more datasets than LDA because it carries less assumptions (Vapnik, 1995, Cristianini and Shawe-Taylor, 2000, Duda et al., 2000, Bishop, 2007). However, the downsides of SVM are the expensive computation with large datasets, the lack of joint distribution, the necessity of complete datasets, and poor performance with unbalanced datasets.

### K-Nearest-Neighbor (KNN)

KNN, as its name indicates, classifies by iteratively assigning each data point to its closest neighbors and then recalculating the center of the class (Duda et al., 2000, Bishop, 2007). KNN does not assume the distribution shape and does not require training. However, it struggles with high dimensional data and unrelated features. The lack of joint distribution and the majority voting mechanisms that the label of a point depends on whichever class has more points closer, also deem KNN to be bad with imbalanced data.

Based on the above descriptions, we predict that when we use the single unit firing to predict the spatial location and valance of the rats in a linear track: 1. The decoding accuracy will increase as the number of neurons being used increases due to the increase in dimensionality. 2. Spatial locations of rats using all cells from the medial temporal lobe can be decoded. 3. Spatial locations of rats can be decoded using neurons at hippocampus and piriform cortex respectively but not amygdala. 4. Linear classifiers (LDA, SVM) will perform better than non-linear classifiers (NB, KNN) in spatial decoding. 5. LDA will perform better than the other classifiers in valence decoding because this classification is imbalanced. 6. SVM, NB and KNN will perform better in classifying the valence of a bin if negative classes are undersampled.

## Methods

### Data Collection

All analyses use previously collected data publicly available and originally published in Girardeau et al. (2017). A summary of the methods used in data recording is provided below.

#### Subjects and electrode implantation

Four individually housed male Long-Evans rats were used in this experiment.Three silicon probes mounted on individual movable microdrives were implanted above the amygdalae bilaterally and in the dorsal hippocampus (refer to Girardeau et al., 2017 for more details). Only two animals (Rat 8 and 9) were selected for our analyses due to missing valence or recording data in the publicly available dataset in the other two animals.

#### Behavioral Paradigm

Animals were trained to run back and forth on a linear track for water rewards (Fig. 1a). In each session an airpuff was presented along the track and was activated only when rats run through one direction. The location and direction of the airpuff changed day-to-day. The position of the animal was tracked using a camera mounted on the ceiling and a red LED attached to the head of the animal (refer to Girardeau et al., 2017 for more details).

**Figure 1.**
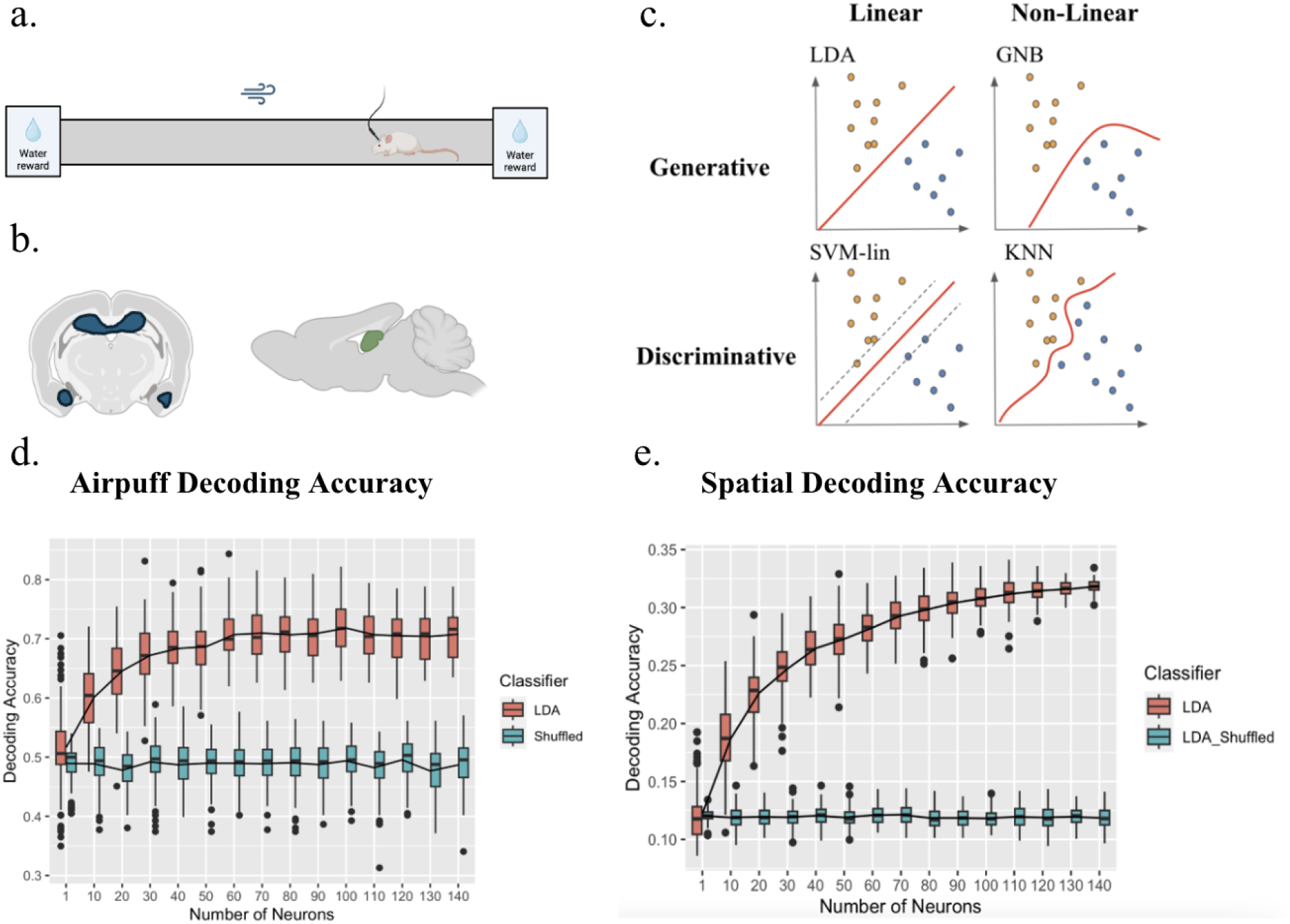
Study Overview. *A. Experiment setup*. A rat is trained to run on a linear track with a water reward on both ends. An airpuff is given at a random location and direction everyday (adapted from Girardeau et al., 2017). *B. Anatomical schematic of recorded regions*. Neurons were recorded in the hippocampus, basolateral amygdala, central amygdala, and piriform cortex. The figure on the left shows the coronal view of these regions, and the figure on the right shows the sagittal view of these regions. *C. Theoretical classifier comparisons*. For this study we compared the results of four popular classifiers: linear discriminant analysis (LDA), which is generative and linear, a linear support vector machine (lin-SVM), which is discriminative and linear, a Gaussian Naive Bayesian classifier (GNB), which is generative and non-linear, and a K-Nearest Neighbors classifier (KNN) which is discriminative and non-linear (adapted from Misaki et al., 2010) *D-E. Decoding accuracy increases as the number of neurons increases*. Panel D shows that the balanced airpuff decoding accuracy increases as the number of neurons included increases before plateauing, for true data using LDA. However, for shuffled data, the accuracy is consistent, indicating the accuracy at chance level 0.5. Panel E shows that the bins decoding accuracy increases before plateauing as the number of neurons included increases for the spatial decoding. However, the shuffled accuracy is consistently at chance level, 0.125, as there are 8 spatial bins. These findings provide guidance for researchers who may be interested in using classification methods with a limited number of recorded neurons.

### Classification Methods

Data was downloaded from https://buzsakilab.nyumc.org/datasets/GirardeauG/ in October, 2022. There are in total 36 sessions, 15 of which contain location, cells spiking and airpuff data, and 9 of which contain more than 100 laps. Thus, these 9 sessions were included for the final analysis.

The linear track was first divided into 10 spatial bins. The beginning and end spatial bin were excluded in the analysis due to, the presence of the water reward, and resulting behavioral variability. The average firing rate within each spatial bin and lap was calculated for each neuron. The bin closest to the airpuff location in each session was defined as the airpuff bin. Due to the limited number of laps, we combined both directional runs of a lap (i.e. left to right and right to left) for classifications. To avoid the bias of imbalanced data, we randomly selected one non-airpuff bin for each iteration.

Neurons from different brain regions are selected based on the labels from the data set (Girardeau et al., 2017). Cells with more than one location label are not included. There are in total 578 neurons in the basolateral amygdala, 156 neurons in central amygdala, and 109 neurons in piriform cortex across the two rats. There are 119 neurons recorded from the hippocampus in one rat.

### Statistical analysis

Analyses were implemented using custom scripts in Matlab 2022b and RStudio version 4.4.1. GraphPad Prism was used to generate figures 2 and 3. All classifiers were implemented in Matlab 2022b using fitc or classify. To allow the classifier to perform the calculations, nan values are converted to zero, and white noise with a mean of 0 and standard deviation of 0.0000001 was added to both training and testing matrix. We utilized leave-one-out cross-validation, in which one lap was excluded in each iteration. For shuffled analyses, training labels were pseudo-randomly assigned to each trial. Classifications were repeated 100 times.

**Figure 2.**
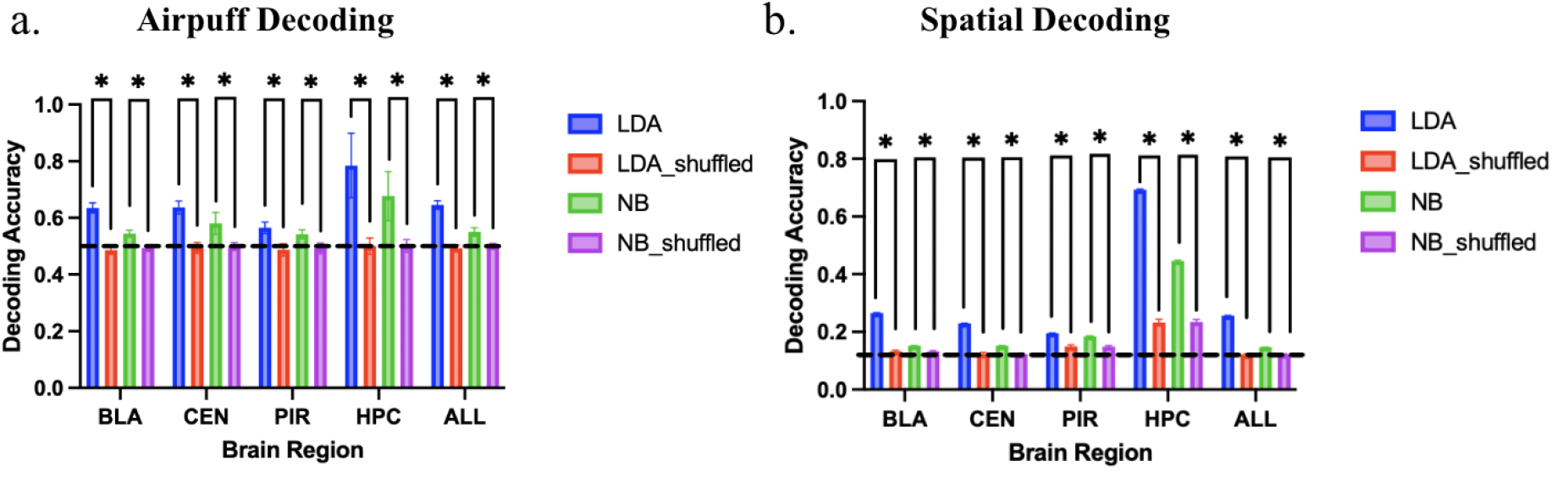
Subregions of the medial temporal lobe can decode space and valence. *A. Airpuff decoding*. The decoding accuracy of balanced airpuff versus non-airpuff decoding is significantly better for true data than shuffled data using all neurons (LDA: p<0.001, NB: p<0.001), and only neurons from basolateral amygdala (LDA: p<0.001, NB: p<0.001), central amygdala (LDA: p<0.001, NB: p<0.001), piriform cortex (LDA: p<0.001, NB: p<0.001), hippocampus (LDA: p<0.001, NB: p<0.001), and using both LDA and NB are significantly better for true data as compared to shuffled (LDA: p=0.002, NB: p=0.013) *B. Spatial Decoding* The decoding accuracy of spatial bins is significantly better for true data than shuffled data from all neurons (LDA: p<0.001, NB: p<0.001), and only neurons from basolateral amygdala (LDA: p<0.001, NB: p<0.001), central amygdala (LDA: p<0.001, NB: p<0.001), piriform cortex (LDA: p<0.001, NB: p<0.001), hippocampus (LDA: p<0.001, NB: p<0.001), and two different classifiers (LDA: p<0.001, NB: p<0.001) chance level 0.125 indicated as dash line).

**Figure 3.**
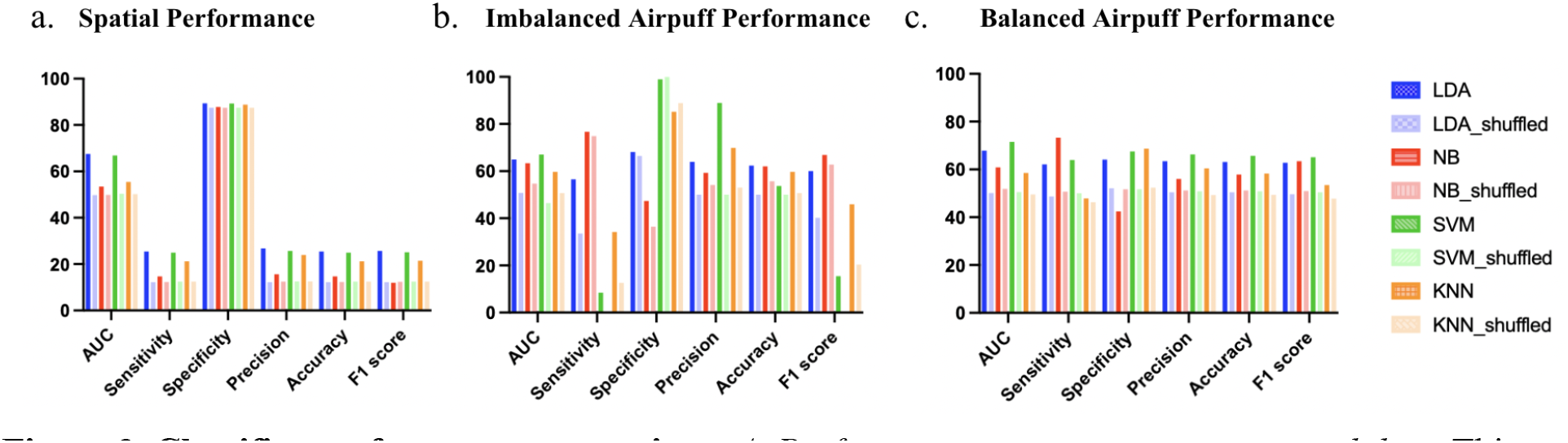
Classifier performance comparison. *A. Performance comparison using spatial data*. This figure compares the performance metrics of spatial decoding (chance accuracy = 0.125) across different classifiers. Linear classifiers LDA and SVM perform the best across metrics. *B. Classifiers performance comparison using imbalanced airpuff data for training*. This figure compares the classifier performance in balanced testing (chance accuracy = 0.5) for the airpuff decoding in imbalanced training such that there are 7 negative classes (non-airpuff bin) but 1 positive class (airpuff bin). NB performs very well across metrics, especially in sensitivity. In contrast, SVM shows near chance accuracy and low sensitivity. *C. Performance comparison using balanced airpuff data for training and testing*. This figure compares the performance in balanced testing (chance accuracy = 0.5) metrics across classifiers when the negative instances (non-airpuff bin) is down-sampled. SVM’s performance has greatly improved after downsampling the majority classes. The irregular high specificity and low sensitivity of SVM and KNN also improve.

#### Spatial decoding

##### Effect of increasing features

We first sought to explore the effect of the number of features on performance accuracy in LDA and NB. To do this, we systematically varied the number of neurons included in the analysis, from 1 to 140, while holding the number of samples constant. 100 iterations were performed at each size, and neurons were randomly selected without replacement for each iteration. Within each iteration, leave-one-lap-out cross validation method was implemented and the accuracy was obtained by averaging the accuracy across laps.

##### Regional comparisons

We first sought to decode if the animal’s spatial location (bin) could be decoded from the neural data. We first implemented LDA and NB using all simultaneously recorded neurons, regardless of regional specification. We then sought to investigate the encoding properties across regions by analyzing neurons from each region separately. LDA and NB are implemented using neurons from the basolateral amygdala (BLA), central amygdala (CEN), hippocampus (HPC), piriform cortex (PIR) separately to classify the animal’s spatial bin. For each session classification, the number of neurons and laps was determined by the number available in that session. The classification is iterated 100 times for each session. Within each iteration, leave-one-lap-out cross validation method is implemented. The final accuracy is obtained by averaging across the lap and sessions. The same classification is also implemented using all neurons as a comparison. A nonparametric Welch t-test was performed between the accuracies of the true data and that of the shuffled data. Bonferroni correction was used for all analyses to correct for multiple comparisons.

##### Classifier comparisons

To compare the performance of different classifiers, the classification was implemented as described above using LDA, NB, lin-SVM, and KNN separately. When applicable, parameters were optimized using 10 fold cross validation and MATLAB’s hyperparameter optimization function. Each session is separately optimized as the number of input features are not the same, parameters are listed in table 2. For baseline comparisons, the parameters are also optimized for the shuffled data (except LDA as it doesn’t have parameters).

**Table 2.** Optimized parameters for each session and each classifier.

| Classifier /Session | 1 | 2 | 3 | 4 | 5 | 6 | 7 | 8 | 9 |
| --- | --- | --- | --- | --- | --- | --- | --- | --- | --- |
| NB | DistributionNames: 'kernel', Width: 2.1023 | DistributionNames: 'kernel', Width: 2.6168 | DistributionNames: 'kernel', Width: 2.8129 | DistributionNames: 'kernel', Width: 2.7248 | DistributionNames: 'kernel', Width: 0.52429 | DistributionNames: 'kernel', Width: 1.2812 | DistributionNames: 'kernel', Width: 0.72271 | DistributionNames: 'kernel', Width: 1.8164 | DistributionNames: 'kernel', Width: 2.565 |
| KNN | NumNeighbors : 21, Distance: 'cityblock' | NumNeighbors : 1, Distance: 'cityblock' | NumNeighbors : 6, Distance: 'cityblock' | NumNeighbors : 54, Distance: 'cityblock' | NumNeighbors : 47, Distance: 'cityblock' | NumNeighbors : 32, Distance: 'cityblock' | NumNeighbors : 20, Distance: 'cityblock' | NumNeighbors : 86, Distance: 'cityblock' | NumNeighbors : 25, Distance: 'cityblock' |
| SVM-lin | Boxconstraint: 0.077023 | Boxconstraint: 0.001 | Boxconstraint: 0.022055 | Boxconstraint: 0.0010011 | Boxconstraint: 0.044751 | Boxconstraint: 0.0052405 | Boxconstraint: 0.0034911 | Boxconstraint: 0.0010025 | Boxconstraint: 0.0025999 |

True positive (TP), true negative (TN), false positive (FP), false negative (FN), and area under the curve (AUC) were calculated for each spatial bin. The metrics are averaged across bins and sessions to create overall performance metrics for each classifier. All results of leave-one-out cross-validation are combined to compute the confusion matrix.

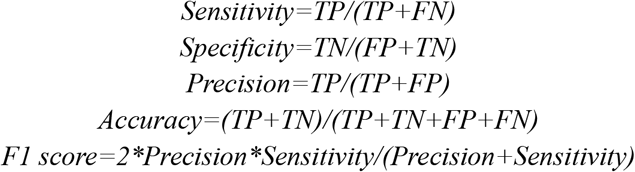

#### Airpuff decoding

##### Region comparison

We sought to determine if the classifiers (LDA and NB) could decode the presence of the airpuff from the neural data. We first used data from all regions in a session that were simultaneously recorded. Similarly, the number of laps and neurons was determined by the number available in each session. We then sought to compare the decoding abilities for the airpuff in the four anatomical regions BLA, CEN, HPC, and PIR. To examine the effect of having balanced data on classifier performance, we downsampled the non-airpuff bin so that only one nonairpuff bin was randomly selected for each iteration to account for the imbalanced dataset (7 nonairpuff bin vs 1 airpuff bin). Each classification was iterated 100 times. Within each iteration, the leave-one-lap-out cross validation method was implemented and the accuracy was obtained by averaging across the lap and sessions. Nonparametric Welch t test was performed between the accuracy of the true data and that of the shuffled data. To correct for multiple comparisons, Bonferroni correction was applied to all analyses.

##### Imbalanced training

To investigate how the classifiers performed with the original imbalanced airpuff data set, we performed the classifier comparison analysis as above, but instead of pseudo-randomly selecting one non-airpuff bin for each lap, we included all bins that do not contain an airpuff for each lap (7), as well as the (1) airpuff bin. To ensure the same chance level performance for fair comparisons between balanced training and imbalanced training, the classifiers were evaluated using a balanced testing set.

## Results

### Decoding accuracy increases with number of features

Increasing the number of neurons increases the classification accuracy (Fig 1d-e). Since the linear discriminant analysis linearly combines all the feature variables, the dimensionality of the function increases as the number of neurons included increases. Thus, the low accuracy when fewer neurons are used indicates at least some level of dimensionality is required for encoding the information, supporting a sparse coding scheme in which many neurons contribute to the information representation. The relative straight line of shuffled data further confirms that when there is no meaningful information contained in the neurons, increasing the number of them doesn’t improve the near-chance accuracy level. The relatively small slope for the Naive Bayesian classifier compared to LDA may be due to its nonlinearity that either overfits the data or deviates from how the information is processed. This may indicate that the information in the neurons are combined in a more linear manner, as linear models show greater increase of accuracy (Glaser et al., 2020, Bishop, 2016).

### Location and Airpuff can be decoded in all brain regions

We then performed classification on neurons individually from the four recorded regions in the medial temporal lobe. As expected, the hippocampus performed significantly better than chance (LDA: p<0.001, NB: p<0.001) and had the highest accuracy in decoding spatial locations (mean accuracy LDA: 0.69, mean accuracy NB: 0.44, chance level: 0.125). However, the other brain regions including piriform cortex and amygdala also performed significantly better than chance levels (fig2b, BLA: LDA: p<0.001, NB: p<0.001; CEN: LDA: p<0.001, NB: p<0.001; PIR: LDA: p<0.001, NB: p<0.001). The result shows that the piriform cortex and amygdala may also contain spatial and valence information. Additionally, the bin containing the airpuff can also be decoded significantly better than shuffle data (BLA: LDA: p<0.001, NB: p<0.001; HPC: LDA: p<0.001, NB: p<0.001; CEN: LDA: p<0.001, NB: p<0.001; PIR: LDA: p<0.001, NB: p<0.001; Across region LDA: p=0.002, NB: p=0.013). In interpreting these results, it is important to consider that the airpuff presents a significant and consistent (within session) spatial marker. This could affect how and why spatial and valence locations are coded across regions.

### Classifier comparisons

#### LDA & SVM

In general, the linear classifiers, LDA and SVM, perform better than the nonlinear classifiers across metrics and decoding tasks (Figure 2, 3). Consistent with our predictions, linear classifiers LDA and SVM performed the best in spatial decoding (Figure 3). While all classifiers showed high specificity scores in spatial decoding, LDA and SVM outperform other classifiers in the rest of the measurements, including AUC, sensitivity, precision, accuracy and F1 score. While LDA performs fairly well in both balanced and imbalanced airpuff decoding, SVM’s performance drops substantially in the imbalanced airpuff decoding. This is manifested as near-chance accuracy, low sensitivity and high specificity, suggesting its accuracy mostly relies on correctly predicting the majority negative instance (non-airpuff). In other words, in an imbalanced dataset where the negative classes are the majority, the classifier will label most instances as negative to achieve high accuracy in training, which will lead to decreased accuracy when it is then presented with balanced testing data.

#### NB

In the spatial decoding task, NB performs the worst across all classifiers. However, in the airpuff decoding, NB performs very well, and is even the best in some metrics like accuracy, F1 score, and sensitivity. Its high performance in sensitivity is particularly obvious in the imbalanced airpuff decoding, making it a potential good candidate for tasks that require a high detection rate for the positive classes and data that is imbalanced.

#### KNN

In the spatial decoding task, KNN’s performance is fairly good, reaching LDA and SVM’s performance in all metrics other than AUC. However, contrary to NB, it performs worst in the airpuff decoding in most metrics. As predicted, KNN is particularly susceptible to the imbalanced data due to its majority voting mechanisms. Though keeping similar accuracy in both balanced and imbalanced data, the increase of specificity and drop of sensitivity suggests that the accuracy comes from overly labeled data to the majority of negative classes. Hence, KNN does not seem to be a good candidate for imbalanced data, and consistently underperforms against other classifiers across measures in this preparation.

In sum, LDA is the most reliable classifier across the three modalities we tested. SVM and KNN are both susceptible to imbalanced data. NB struggles in the spatial decoding task, but is particularly good at correctly labeling the positive instances even in imbalanced data.

## Discussion

Aligned with our hypothesis, we found that the spatial location can be decoded using all recorded cells from the medial temporal lobe, cells from hippocampus, and cells from piriform cortex. Unexpectedly, we found that space can also be decoded in amygdala, suggesting a potential distributed encoding of space across medial temporal lobe. The decoding accuracy increases as the number of neurons included increases. The classifiers’ performance also follows our prediction in that the linear classification performs the best overall, particularly in spatial decoding. In the imbalanced valence decoding, NB outperforms other classifiers overall, as demonstrated by the highest composite F1 score. In the balanced valence decoding, there is not a classifier that is best in all performance metrics. While NB has the highest sensitivity, SVM has the highest accuracy, F1 score and area under curve. Under sampling of the imbalanced class improved the performance of SVM and KNN, but not LDA and NB. Our study shows an unknown advantage of NB in imbalanced data given its high performance in airpuff decoding regardless of sampling methods. These results further demonstrate that there is not a global optimal classifier but local optimal classifier for each task.

### The Amygdala encodes space

Previous work investigating the role of the amygdala in spatial navigation has shown that it is not essential for navigation to occur (Peinado-Manzano, 1990). However, the amygdala does appear to play some role in facilitating the hippocampus’s encoding of spatial information in stimulation and lesion studies (Yang et al., 2016, Yang & Wang, 2017, Inman et al., 2023). Yet, prior work has not investigated whether the spatial information can be decoded independently in amygdala. Our results expand on the nuanced role the amygdala may play in spatial coding in the intact brain by showing that spatial information can be decoded in the amygdala. It is possible that the decodability is due to its anatomical and functional connections with the hippocampus. It is important to consider the role of the task in influencing the firing patterns in each region. We may see increased spatial representations in the amygdala due to the presence of high valence locations (reward ports and airpuff location). However, our findings suggest that the amygdala is capable of carrying decodable spatial representations. Hence, future studies are needed to further investigate the role of task demands and inherent capacity of the amygdala in spatial encoding.

### Linear classifiers outperform nonlinear classifiers

Additionally, this study provides important insights for choosing a classifier when performing classification or prediction tasks using single cell unit data. Linear classifiers like LDA and SVM are easy to implement and have high performance, as measured by Accuracy, Specificity, Sensitivity, F1 score and AUC. However, the NB decoder outperforms other classifiers with imbalanced data, particularly in accurately identifying positive instances.These analyses further demonstrate the importance of considering the most important performance metric for the classifier given data structure and study goals, particularly for imbalanced data.

### Limitations

One consideration of this study in comparing results to other linear track experiments, is the presence of the aversive air puff. This additional variable may boost decoding by providing a highly valentanchor for the spatial maps, increasing classification accuracy. This may be particularly relevant for the amygdala’s spatial decoding performance, as it is related to danger processing. The linear track also limits rats’ movement in the one dimensional track which may increase the spatial information by providing boundary anchor points, as well as greatly reducing the dimension and complexity of spatial information (Yang et al., 2024). It is possible that when rats are allowed to explore bigger regions without a negative stimulus, the decoding performance of amygdala may drop, as it is likely not specialized for spatial processing with high granularity.

In order to facilitate comparisons across the imbalanced and balanced binary airpuff classifications, all classifiers were tested with balanced testing data sets. This aids in the comparison of accuracies, because chance is held constant across the two sets. However, this penalizes classifiers, such as the Naive Bayesian and K-Nearest Neighbors, which account for group probabilities when determining classifications. Future work should include comparisons of classifiers trained and tested on imbalanced data.

In sum, this paper uses different classifiers to investigate whether space and valence can be decoded using the neural firing data in hippocampus, the amygdala and the piriform cortex. It adds to the understanding of how spatial information is distributed in the medial temporal lobe by highlighting the ability to decode spatial information in the amygdala and piriform cortex. This paper also highlights the strengths and weaknesses of different classifiers, and illustrates the effects of classifier choice on results. These analyses can be used to help guide investigator’s choices when analyzing their own datasets.

## References

Astrand, E., Enel, P., Guilhem Ibos, Peter Ford Dominey, Baraduc, P., & Suliann Ben Hamed. (2014). Comparison of Classifiers for Decoding Sensory and Cognitive Information from Prefrontal Neuronal Populations. PloS One, 9(1), e86314–e86314. 10.1371/journal.pone.0086314

Bekkers, J. M., & Suzuki, N. (2013). Neurons and circuits for odor processing in the piriform cortex. Trends in Neurosciences, 36(7), 429–438. 10.1016/j.tins.2013.04.005

Bird, C. M., & Burgess, N. (2008). The hippocampus and memory: insights from spatial processing. Nature Reviews. Neuroscience, 9(3), 182–194. 10.1038/nrn2335

Bishop, C. M. (1993). Novelty Detection and Neural Network Validation. Springer EBooks, 789–794. 10.1007/978-1-4471-2063-6_225

Bishop, C. M. (2006). Pattern Recognition and Machine Learning (pp. 43–44). Springer.

Bishop, C. M. (2016). Pattern Recognition and Machine Learning. SpringerLink. 10.1007-978-0-387-45528-0

Blazing, R. M., & Franks, K. M. (2020). Odor coding in piriform cortex: mechanistic insights into distributed coding. Current Opinion in Neurobiology, 64, 96–102. 10.1016/j.conb.2020.03.001

Burgess, N., Maguire, E. A., & O’Keefe, J. (2002). The Human Hippocampus and Spatial and Episodic Memory. Neuron, 35(4), 625–641. 10.1016/s0896-6273(02)00830-9

Edvard Ingjald Moser, Moser, M.-B., & McNaughton, B. (2017). Spatial representation in the hippocampal formation: a history. Nature Neuroscience, 20(11), 1448–1464. 10.1038/nn.4653

Eichenbaum, H. (2017). Prefrontal–hippocampal interactions in episodic memory. Nature Reviews. Neuroscience, 18(9), 547–558. 10.1038/nrn.2017.74

Girardeau, G., Inema, I., & György Buzsáki. (2017). Reactivations of emotional memory in the hippocampus–amygdala system during sleep. Nature Neuroscience (Print), 20(11), 1634–1642. 10.1038/nn.4637

Glaser, J. I., Benjamin, A. S., Chowdhury, R. H., Perich, M. G., Miller, L. E., & Kording, K. P. (2020). Machine learning for neural decoding. Eneuro, ENEURO.0506-19.2020. 10.1523/eneuro.0506-19.2020

Grijseels, D. M., Shaw, K., Barry, C., & Hall, C. N. (2021). Choice of method of place cell classification determines the population of cells identified. PLOS Computational Biology/PLoS Computational Biology, 17(7), e1008835–e1008835. 10.1371/journal.pcbi.1008835

Haxby, J. V., M. Ida Gobbini, Furey, M. L., Alumit Ishai, Schouten, J. L., & Pietrini, P. (2001). Distributed and Overlapping Representations of Faces and Objects in Ventral Temporal Cortex. Science, 293(5539), 2425–2430. 10.1126/science.1063736

Haynes, J.-D., & Rees, G. (2006). Decoding mental states from brain activity in humans. Nature Reviews Neuroscience, 7(7), 523–534. 10.1038/nrn1931

Herry, C., Ciocchi, S., Senn, V., Demmou, L., Müller, C., & Lüthi, A. (2008). Switching on and off fear by distinct neuronal circuits. Nature, 454(7204), 600–606. 10.1038/nature07166

Inman, C. S., Hollearn, M. K., Augustin, L., Campbell, J. M., Olson, K. L., & Wahlstrom, K. L. (2023). Discovering how the amygdala shapes human behavior: From lesion studies to neuromodulation. Neuron, 111(24), 3906–3910. 10.1016/j.neuron.2023.09.040

J. O’Keefe, & J. Dostrovsky. (1971). The hippocampus as a spatial map. Preliminary evidence from unit activity in the freely-moving rat. Brain Research, 34(1), 171–175. 10.1016/0006-8993(71)90358-1

Johannes Niediek, & Bain, J. (2014). Human single-unit recordings reveal a link between place-cells and episodic memory. Frontiers in Systems Neuroscience, 8. 10.3389/fnsys.2014.00158

Kim, T., & Lee, J.-S. (2023). Maximizing AUC to learn weighted naive Bayes for imbalanced data classification. Expert Systems with Applications, 217, 119564–119564. 10.1016/j.eswa.2023.119564

Kong, M.-S., Eun Joo Kim, Park, S., Zweifel, L. S., Huh, Y., Cho, J., & Kim, J. J. (2021, September 17). “Fearful-place” coding in the amygdala-hippocampal network. ELife; eLife Sciences Publications, Ltd. https://elifesciences.org/articles/72040#s4

LeDoux, J. (2007). The amygdala. Current Biology, 17(20), R868–R874. 10.1016/j.cub.2007.08.005

Mandelkow, H., de, A., & Duyn, J. H. (2016). Linear Discriminant Analysis Achieves High Classification Accuracy for the BOLD fMRI Response to Naturalistic Movie Stimuli. Frontiers in Human Neuroscience, 10. 10.3389/fnhum.2016.00128

Maren, S. (2001). Neurobiology of Pavlovian Fear Conditioning. Annual Review of Neuroscience, 24(1), 897–931. 10.1146/annurev.neuro.24.1.897

Masaya Misaki, Kim, Y., Bandettini, P. A., & Nikolaus Kriegeskorte. (2010a). Comparison of multivariate classifiers and response normalizations for pattern-information fMRI. NeuroImage, 53(1), 103–118. 10.1016/j.neuroimage.2010.05.051

Masaya Misaki, Kim, Y., Bandettini, P. A., & Nikolaus Kriegeskorte. (2010b). Comparison of multivariate classifiers and response normalizations for pattern-information fMRI. NeuroImage, 53(1), 103–118. 10.1016/j.neuroimage.2010.05.051

Mehrad Sarmashghi, Jadhav, S. P., & Eden, U. T. (2022). Integrating Statistical and Machine Learning Approaches for Neural Classification. IEEE Access, 10, 119106–119118. 10.1109/access.2022.3221436

Mitchell, T. M., Hutchinson, R., Just, M. A., Niculescu, R. S., Pereira, F., & Wang, X. (2003). Classifying instantaneous cognitive states from FMRI data. AMIA … Annual Symposium Proceedings. AMIA Symposium, 2003, 465–469. https://www.ncbi.nlm.nih.gov/pmc/articles/PMC1479944/

Moser, E. I., Moser, M.-B., & McNaughton, B. L. (2017). Spatial representation in the hippocampal formation: a history. Nature Neuroscience, 20(11), 1448–1464. 10.1038/nn.4653

O’Keefe, J. (1976). Place units in the hippocampus of the freely moving rat. Experimental Neurology, 51(1), 78–109. 10.1016/0014-4886(76)90055-8

Olga Therese Ousdal, Specht, K., Server, A., Andreassen, O. A., Dolan, R. J., & Jensen, J. (2014). The human amygdala encodes value and space during decision making. NeuroImage, 101, 712–719. 10.1016/j.neuroimage.2014.07.055

Park, D. J., Park, M. W., Lee, H., Kim, Y.-J., Kim, Y., & Park, Y. H. (2021). Development of machine learning model for diagnostic disease prediction based on laboratory tests. Scientific Reports, 11(1). 10.1038/s41598-021-87171-5

Peck, C. J., & C Daniel Salzman. (2014). Amygdala neural activity reflects spatial attention towards stimuli promising reward or threatening punishment. ELife, 3. 10.7554/elife.04478

Peck, C. J., Lau, B., & Salzman, C. D. (2013). The primate amygdala combines information about space and value. Nature Neuroscience, 16(3), 340–348. 10.1038/nn.3328

Peinado-Manzano, M. A. (1990). The role of the amygdala and the hippocampus in working memory for spatial and non-spatial information. Behavioural Brain Research, 38(2), 117–134. 10.1016/0166-4328(90)90010-c

Pinelopi Kyriazi, Headley, D. B., & Pare, D. (2018). Multi-dimensional Coding by Basolateral Amygdala Neurons. Neuron, 99(6), 1315-1328.e5. 10.1016/j.neuron.2018.07.036

Poo, C., Agarwal, G., Niccolò Bonacchi & Mainen, Z. F. (2021). Spatial maps in piriform cortex during olfactory navigation. Nature, 601(7894), 595–599. 10.1038/s41586-021-04242-3

Qu, L., & Pei, Y. (2024). A Comprehensive Review on Discriminant Analysis for Addressing Challenges of Class-Level Limitations, Small Sample Size, and Robustness. Processes, 12(7), 1382. 10.3390/pr12071382

R. Adolphs, Tranel, D., H. Damasio, & A. Damasio. (1994). Impaired recognition of emotion in facial expressions following bilateral damage to the human amygdala. Nature, 372(6507), 669–672. 10.1038/372669a0

Rayan Alanazi. (2022). Identification and Prediction of Chronic Diseases Using Machine Learning Approach. Journal of Healthcare Engineering, 2022, 1–9. 10.1155/2022/2826127

Rehan Akbani, Kwek, S., & Japkowicz, N. (2004). Applying Support Vector Machines to Imbalanced Datasets. Lecture Notes in Computer Science, 39–50. 10.1007/978-3-540-30115-8_7

Rigotti, M., Barak, O., Warden, M. R., Xiao Jing Wang, Daw, N. D., Miller, E. K., & Fusi, S. (2013). The importance of mixed selectivity in complex cognitive tasks. Nature (London), 497(7451), 585–590. 10.1038/nature12160

Scholz, M., & Wimmer, T. (2021). A comparison of classification methods across different data complexity scenarios and datasets. Expert Systems with Applications, 168, 114217–114217. 10.1016/j.eswa.2020.114217

Shinde, A., Mohapatra, S., & Gottfried Schlaug. (2023). Identifying the engagement of a brain network during a targeted tDCS-fMRI experiment using a machine learning approach. PLoS Computational Biology, 19(4), e1011012–e1011012. 10.1371/journal.pcbi.1011012

Stefanini, F., Kushnir, L., Jimenez, J. C., Jennings, J. H., Woods, N. I., Stuber, G. D., Kheirbek, M. A., Hen, R., & Fusi, S. (2020). A Distributed Neural Code in the Dentate Gyrus and in CA1. Neuron, 107(4), 703-716.e4. 10.1016/j.neuron.2020.05.022

Uddin, S., Khan, A., Md Ekramul Hossain, & Mohammad Ali Moni. (2019). Comparing different supervised machine learning algorithms for disease prediction. BMC Medical Informatics and Decision Making, 19(1). 10.1186/s12911-019-1004-8

Wixted, J. T., Squire, L. R., Jang, Y., Papesh, M. H., Goldinger, S. D., Kuhn, J. R., Smith, K. A., Treiman, D. M., & Steinmetz, P. N. (2014). Sparse and distributed coding of episodic memory in neurons of the human hippocampus. Proceedings of the National Academy of Sciences of the United States of America, 111(26), 9621–9626. 10.1073/pnas.1408365111

Wolpert, D. H., & Macready, W. G. (1997). No free lunch theorems for optimization. IEEE Transactions on Evolutionary Computation, 1(1), 67–82. 10.1109/4235.585893

Wood, E. R., Dudchenko, P. A., & Eichenbaum, H. (1999). The global record of memory in hippocampal neuronal activity. Nature, 397(6720), 613–616. 10.1038/17605

Xie, J., & Qiu, Z. (2007). The effect of imbalanced data sets on LDA: A theoretical and empirical analysis. Pattern Recognition, 40(2), 557–562. 10.1016/j.patcog.2006.01.009

Xue, J.-H., & Titterington, D. M. (2008). Do unbalanced data have a negative effect on LDA? Pattern Recognition, 41(5), 1558–1571. 10.1016/j.patcog.2007.11.008

Yang, X., Cacucci, F., Burgess, N., Wills, T. J., & Chen, G. (2024). Visual boundary cues suffice to anchor place and grid cells in virtual reality. Current Biology, 34(10), 2256-2264.e3. 10.1016/j.cub.2024.04.026

Yang, Y., & Wang, J.-Z. (2017). From Structure to Behavior in Basolateral Amygdala-Hippocampus Circuits. Frontiers in Neural Circuits, 11. 10.3389/fncir.2017.00086

Yang, Y., Wang, Z.-H., Jin, S., Gao, D., Liu, N., Chen, S.-P., Zhang, S., Liu, Q., Liu, E., Wang, X., Liang, X., Wei, P., Li, X., Li, Y., Yue, C., Li, H., Wang, Y.-L., Wang, Q., Ke, D., & Xie, Q. (2016). Opposite monosynaptic scaling of BLP–vCA1 inputs governs hopefulness- and helplessness-modulated spatial learning and memory. Nature Communications, 7(1). 10.1038/ncomms11935

Yang, Z., Huang, Z., Gonzalez-Castillo, J., Dai, R., Georg Northoff, & Bandettini, P. (2014). Using fMRI to decode true thoughts independent of intention to conceal. NeuroImage, 99, 80–92. 10.1016/j.neuroimage.2014.05.034

